# The *C. elegans* NF2/Merlin Molecule NFM-1 Non-Autonomously Regulates Neuroblast Migration and Interacts Genetically with the Guidance Cue SLT-1/Slit

**DOI:** 10.1101/055079

**Authors:** Matthew P. Josephson, Rana Aliani, Erik Lundquist

## Abstract

During nervous system development, neurons and their progenitors often migrate to their final destinations. In *Caenorhabditis elegans*, the bilateral Q neuroblasts and their descendants migrate long distances in opposite directions, despite being born in the same posterior region. QR on the right migrates anteriorly and generates the AQR neuron positioned near the head, and QL on the left migrates posteriorly, giving rise to the PQR neuron positioned near the tail. In a screen for genes required for AQR and PQR migration, we identified an allele of *nfm-1*, which encodes a molecule similar to vertebrate NF2/Merlin, an important tumor suppressor in humans. Mutations in *NF2* lead to Neurofibromatosis Type II, characterized by benign tumors of glial tissues. These molecules contain <u>F</u>our-point-one <u>E</u>zrin <u>R</u>adixin <u>M</u>oesin (FERM) domains characteristic of cytoskeletal-membrane linkers, and vertebrate NF2 is required for epidermal integrity. Vertebrate NF2 can also regulate several transcriptional pathways including the Hippo pathway. Here we demonstrate that in *C. elegans*, *nfm-1* is required for complete migration of AQR and PQR, and that it likely acts outside of the Q cells themselves in a non-autonomous fashion. We also show a genetic interaction between *nfm-1* and the *C. elegans Slit* homolog *slt-1*, which encodes a conserved secreted guidance cue. In vertebrates, *NF2* can control *Slit2* mRNA levels through the hippo pathway in axon pathfinding, suggesting a conserved interaction of NF2 and Slit2 in regulating migration.

## Introduction

A critical process in nervous system development is the directed migration of neurons to precise destinations. Directed migration is a complex process that requires integration of extracellular cues into cytoskeletal changes which guide the cell to a specific location. In *C. elegans* the Q neuroblasts are an established system to study directed cell migrations (MIDDELKOOP AND KORSWAGEN 2014). The QR and QL neuroblasts are born in the posterior region of the worm yet migrate in opposite directions (SULSTON AND HORVITZ 1977; SALSER AND KENYON 1992; SALSER *et al.* 1993). QL is born on the left side of the animal and migrates posteriorly over the seam cell V5 before dividing. During this initial migration QL detects a posteriorly derived EGL-20/Wnt signal which through canonical Wnt signaling initiates transcription of *mab-5/Hox* (SALSER AND KENYON 1992). MAB-5 drives further posterior migration of the QL lineage, resulting in the QL.a descendant PQR migrating to the tail near the anus and posterior phasmid ganglion. QR is born on the right side of the animal and migrates anteriorly over the seam cell V4 and away from the EGL-20/Wnt signal (SALSER *et al.* 1993; HARRIS *et al.* 1996; SALSER AND KENYON 1996). QR does not initiate *mab-5* expression in response to Wnt and continues to migrate anteriorly. After division, QR.a undergoes an identical pattern of cell divisions and cell death as QL.a while migrating anteriorly, with AQR completing migration near the posterior pharyngeal bulb in the head (Figure 1)(MALOOF *et al.* 1999; WHANGBO AND KENYON 1999). Initial Q migrations are controlled autonomously by the receptor molecules UNC-40/DCC and PTP-3/LAR (HONIGBERG AND KENYON 2000; SUNDARARAJAN AND LUNDQUIST 2012) and non-autonomously by the Fat-like cadherin CDH-4 (SUNDARARAJAN *et al.* 2014). Later Q descendant migrations are controlled by Wnt signaling (WHANGBO AND KENYON 1999; ZINOVYEVA AND FORRESTER 2005; ZINOVYEVA *et al.* 2008; HARTERINK *et al.* 2011), which appears to not be involved in initial migration (JOSEPHSON *et al.* 2016), and by the transmembrane receptor MIG-13 in parallel with SDN-1/Syndecan (WANG *et al.* 2013; SUNDARARAJAN *et al.* 2015).

**Figure 1.**
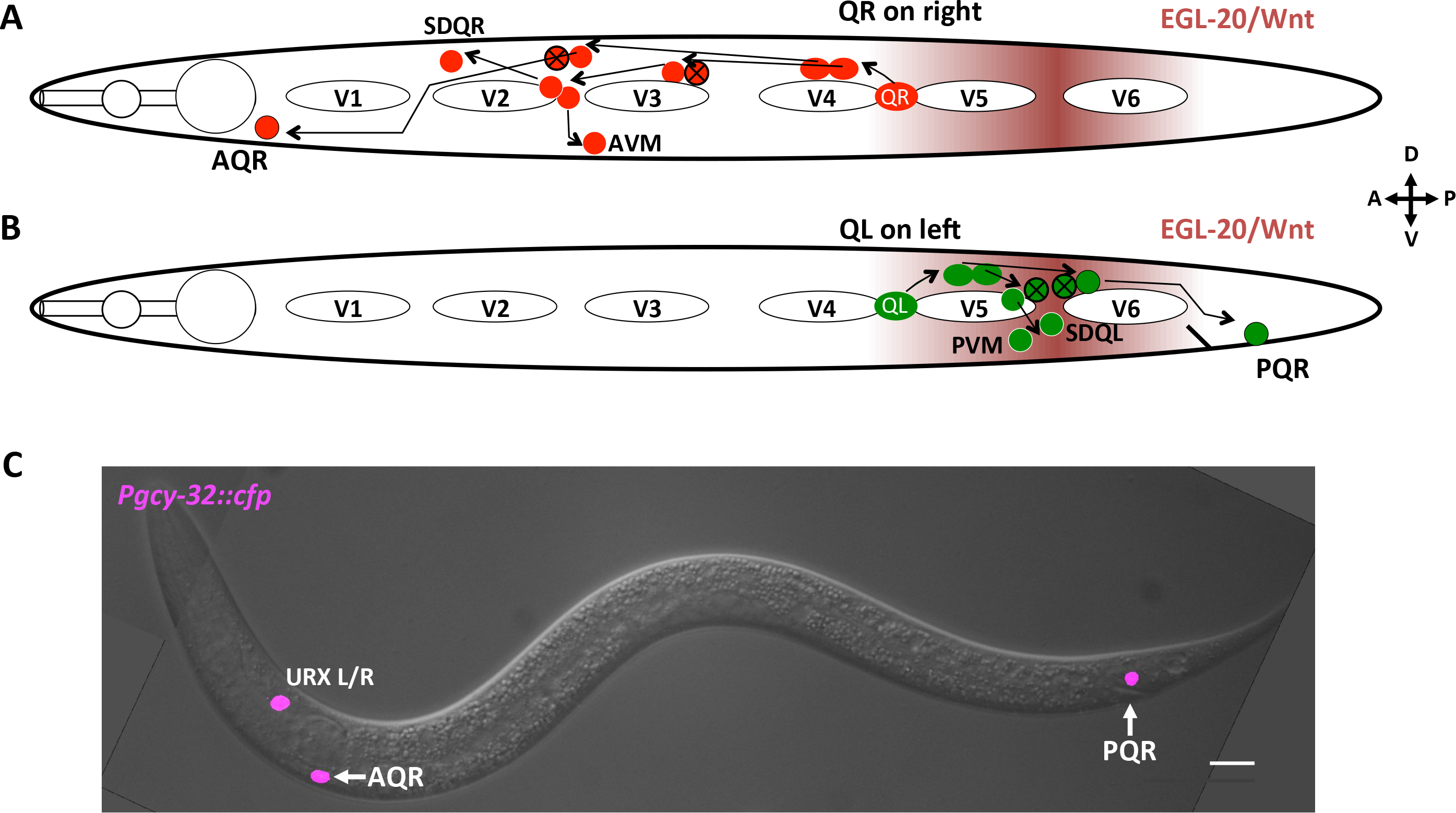
Migration of QR and QL descendants. A-B) Diagrams representing the migration and cell division pattern of QR on the right side (A) and QL on the left side (B) in the L1 animal, showing birthplace of the Q neuroblasts, and approximate locations of the Q descendants. Maroon shading represents the posteriorly derived EGL-20/Wnt signal. White ovals are hypodermal seam cells V1-V6. Circles with black x indicate cells that undergo programmed cell death after cell division. Dorsal is up, anterior to the left. C) Merged DIC and fluorescent micrograph showing location of Q descendants AQR and PQR in an adult wild-type animal. *Pgcy-32::cfp* is expressed in AQR, PQR and URXL/R. The scale bar represents 10μM

To identify additional molecules that regulate Q migrations, a forward genetic screen for mutations affecting AQR and PQR migration was previously performed. Here we report that this screen identified an allele of *nfm-1*, which encodes a *C. elegans* Neurofibromatosis Type II (NF2)/Merlin molecule. *NF2* acts as a tumor suppressor in humans, and mutations in the gene lead to development of neurofibromatosis type II (GUSELLA *et al.* 1996; GUTMANN *et al.* 1997), a disease of benign Schwannomas that must be removed surgically. NF2/Merlin is involved in signaling pathways involving hippo, mTOR and PI3K-Akt (ZHAO *et al.* 2007; STRIEDINGER *et al.* 2008; JAMES *et al.* 2009; OKADA *et al.* 2009). Additionally, NF2 is involved in nervous system maintenance, corpus callosum development, and axon guidance (SCHULZ *et al.* 2013; LAVADO *et al.* 2014; SCHULZ *et al.* 2014). In corpus callosum development, NF2 inhibits the hippo pathway component Yap. In *NF2* mutants, this inhibition is relieved, resulting in increased expression of *Slit2*, a secreted axon guidance cue that prevents midline crossing. This leads to defects in midline crossing of axons in the callosum (LAVADO *et al.* 2014).

In *C. elegans*, RNAi against *nfm-1* results in embryonic lethality (SKOP *et al.* 2004), and an *nfm-1::gfp* transgene is reported as being localized to the basolateral region of gut epithelium (ZHANG *et al.* 2011). Here we report that two likely hypomorphic mutations in *nfm-1* display AQR and PQR migration defects. Mosaic analysis and expression studies indicated that NFM-1 might not act in the Q cells themselves (i.e. that it acts non-autonomously). Finally, we show a genetic interaction between NFM-1 and the secreted guidance cue SLT-1 in AQR migration. In vertebrates, Slit1 and Slit2 are required for guidance of many axons, acting through the Robo family receptors (NGUYEN BA-CHARVET *et al.* 1999; PIPER *et al.* 2000; BAGRI *et al.* 2002; UNNI *et al.* 2012; KIM *et al.* 2014). The Slit-Robo guidance pathway is conserved in *C. elegans* where SLT-1 acts as a guidance cue for several neurons through SAX-3/Robo (HAO *et al.* 2001; CHANG *et al.* 2006; QUINN *et al.* 2006; XU AND QUINN 2012). In general, detection of extracellular guidance cues such as *Slit* cause cytoskeletal changes that result in directed migration of cells and axonal growth cones, most typically repulsion. We show that *slt-1* mutations enhance AQR migration defects of *nfm-1* mutations, and that *sax-3* mutants display defects in AQR and PQR migration. These results are consistent with a model in which NFM-1 regulates AQR and PQR migration by controlling the production of an extracellular cue, either SLT-1 itself or an unidentified cue that acts in parallel to SLT-1.

## Materials and Methods

### Nematode Strains and Genetics

*C. elegans* were grown under standard conditions at 20°C on Nematode Growth Media (NGM) plates (SULSTON AND BRENNER 1974). N2 Bristol was the wild-type strain. Alleles used include LG III: *nfm-1(ok754), nfm-1(lq132)*, LG X: *sax-3(ky123), slt-1(ok255), slt-1(eh15)*. Standard gonadal injection was used to create the following extrachromosomal arrays: *lqEx773[nfm-1::gfp* fosmid (5ng/μL), *Pgcy::32::yfp* (50ng/μL)], *lqEx782* [*Pnfm-1::gfp* (10ng/μL), *Pgcy-32::cfp* (25ng/μL)]. Ultraviolet Trimethylpsoralen (UV/TMP) techniques (MELLO AND FIRE 1995) were used to integrate extrachromosomal arrays to generate the following transgenes: LGII: *lqIs244 [lqEx737, Pgcy-32::cfp* (25ng/μL)], unknown chromosomal location *lqIs247 [lqEx773, nfm-1::gfp], lqIs274 [lqEx834, Pegl-17::myr-mCherry* (20ng/μL) *Pegl-l7::mCherry::his-24* (20ng/μL)]. The *nfm-1::gfp* fosmid was obtained from the TransgeneOme project, clone 7039520022144752 D02 (SAROV *et al.* 2006). *nfm-1(ok754)* was maintained as a heterozygote over the *hT2* balancer because homozygous *ok754* animals arrest during larval stages, but positions of AQR and PQR could be scored in these arrested animals. Genotypes with M+ had maternal contribution from the *hT2* balancer.

### Scoring migration of AQR and PQR

AQR migrates to a position near post-deirid ganglia in the region of the posterior pharyngeal bulb, and PQR migrates posteriorly to a position near the phasmid ganglia posterior to the anus. We used a method as described previously to score AQR and PQR position using *Pgcy-32* to drive expression of fluorescent proteins (SHAKIR *et al.* 2006; CHAPMAN *et al.* 2008). Five positions in the anterior-posterior axis of the animal were used to score AQR and PQR position. Position 1 was the wild-type position of AQR and is the region around the posterior pharyngeal bulb. Neurons anterior to the posterior pharyngeal bulb were not observed. Position 2 was posterior to position 1, but anterior to the vulva. Position 3 was the region around the vulva, position 4 was the birthplace of Q cells, and position 5 was posterior to the anus, the wild-type position of PQR (see Figure 2D). A Leica DM550 equipped with YFP, CFP, GFP, and mCherry filters, was used to acquire all micrographs, and for visualization of A/PQR for scoring. Micrographs were acquired using a Qimaging Retiga camera. Significances of difference were determined using Fisher’s Exact test.

**Figure 2.**
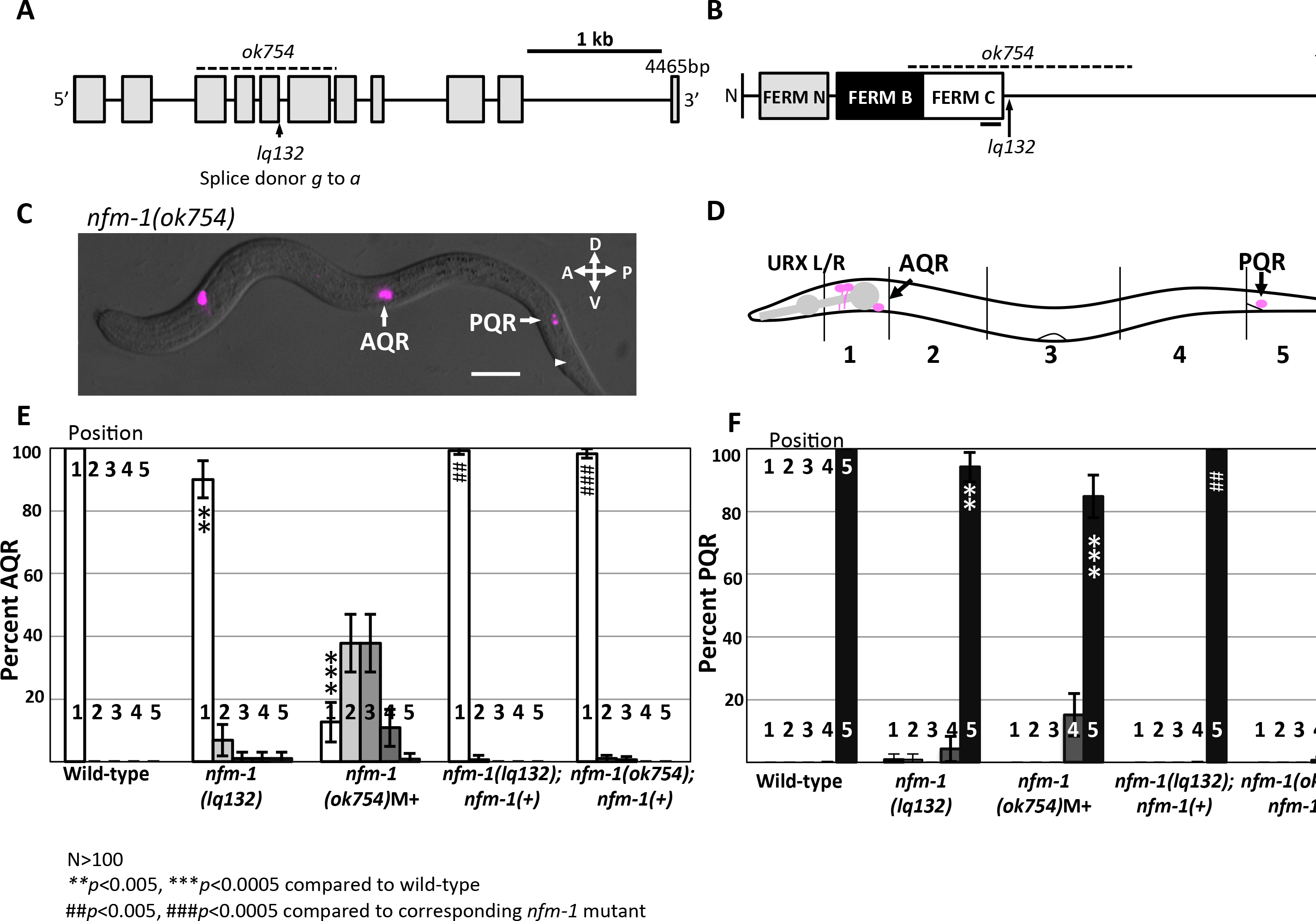
Position of Q descendants AQR and PQR in *nfm-1* mutants. A) A diagram of the *nfm-1* locus and alleles used. The *ok754* deletion (dashed line), and *lq132* splice site mutation (arrow) are noted. B) NFM-1 isoform A domain structure and allele locations are shown. The FERM domain lobes N (gray), B (black), and C (white) are shown. The black bar under FERM C represents predicted actin-binding motif. The dashed line is *ok754* in frame deletion, and *lq132* splice donor mutation location is marked by an arrow. C) Merged DIC and fluorescent micrograph of an *nfm-1(ok754)* arrested larval mutant animal. Both AQR and PQR failed to migrate (PQR wild-type position noted by arrowhead). Scale bar represents 10μM. D) Diagram of scoring positions in an L4 animal, with wild-type locations of AQR and PQR shown as magenta circles. E-F) Chart showing percent of AQR (E) or PQR (F) in positions 1-5 in different genotypes. All animals unless otherwise noted were scored using *lqIs58 (Pgcy-32::cfp).* M+ indicates animals were scored from heterozygous mother and have wild-type maternal contribution of *nfm-1. nfm-1(+)* animals harbor the array containing the *nfm-1::gfp* fosmid. Asterisks indicate significant) difference from wild-type (N>100 *\*p*<0.05, \*\**p*<0.005, \*\*\**p*<0.0005 Fisher’s exact test). Pound signs indicate, for that position, a significant rescue of corresponding *nfm-1* mutant (N>100 #*p*0.05, ##*p*<0.005, ###*p*<0.0005, Fisher’s exact test). Error bars represent two times the standard error of the proportion.

### Mosaic analysis

Mosaic analysis was conducted as previously described (CHAPMAN *et al.* 2008; SUNDARARAJAN *et al.* 2014). A rescuing *nfm-1(+)* extrachromosomal array was generated using the *nfm-1::gfp* fosmid with a *Pgcy-32::yfp* marker. This array was crossed into *nfm-1(ok754)/hT2; lqIs58 (Pgcy-32::cfp)* to create the rescuing array *lqEx773*, referred to as *nfm-1(+)*. Presence of the rescuing array was determined by *Pgcy-32::yfp* expression, and position of AQR and PQR was determined by *Pgcy-32::cfp* expression. *nfm-1(ok754)IU; nfm-1(+)* animals were viable and fertile and had wild-type AQR and PQR position, indicating rescue of *nfm-1(ok754)*. Presence of YFP in AQR or PQR indicated *nfm-1(+)* was present in those cells during their migrations. *Pgcy-32* is also expressed in URX, and presence of YFP in the URX neurons indicates other tissues have inherited *nfm-1(+)*. Animals that lost *nfm-1(+)* in AQR or PQR, and retained *nfm-1(+)* in the other Q descendant (PQR and AQR respectively) and URX were scored for AQR and PQR position.

### Synchronization of L1 larvae for expression analysis

L1 Animals carrying *Pnfm-1::gfp* or *nfm-1::gfp* fosmid were synchronized as described previously to the time of Q cell migration (3-5h post hatching)(CHAPMAN *et al.* 2008; SUNDARARAJAN AND LUNDQUIST 2012). Gravid adults were allowed to lay eggs overnight. Plates were washed with M9 buffer, and eggs remained attached to plate. Hatched larvae were collected every half hour using M9 washes and placed onto clean NGM plates for later imaging. *Pegl-17::mCherry* was used as a Q cell marker to determine overlapping expression of *nfm-1* expression constructs.

## Results

### *nfm-1* mutants have defective AQR and PQR migration

QL and QR undergo identical patterns of cell division, cell death, and neuronal differentiation, but migrate in opposite directions (Figure 1). To identify genes required for AQR and PQR migration, a forward genetic screen using the mutagen ethyl methanesulfonate (EMS) was conducted (SULSTON AND HODGKIN 1988). This screen identified a new mutation with defective AQR and PQR migration, *lq132* (10% AQR defects and 6% PQR defects, Figure 2). The genome of the *lq132*-bearing strain was sequenced and variants were detected using Cloudmap (MINEVICH *et al.* 2012). The strain contained a contained a splice-donor mutation after the fifth exon in the *nfm-1* gene (Figure 2A) (<u>G</u>TATGTGT to <u>A</u>TATGTGT). We scored AQR and PQR migration in the *nfm-1(ok754)* mutant generated by the *C. elegans* gene knock-out consortium. *nfm-1(ok754)* is an in-frame 1042-bp deletion with breakpoints in exons 3 and 7 that removes all of exons 4, 5, and 6 (Figure 2A and Figure 2B). *nfm-1(ok754)* homozygotes arrested as larvae, but we were able to score AQR and PQR position in arrested larvae. *nfm-1(ok754)* had strong AQR defects, with 88% of AQR failing to migrate to the head, and occasional (1%) posterior AQR migration (Figure 2C and Figure 2E). *nfm-1(ok754)* also had significant PQR defects, with 15% of PQR failing to migrate into the wild-type position 5 posterior to the anus (Figure 2F). An *nfm-1::gfp* fosmid transgene rescued AQR and PQR defects of both *lq132* and *ok754*, indicating that mutations in *nfm-1* were causative for the migration deficiencies of *lq132* and *ok754* (Figure 2E and Figure 2F).

*nfm-1* encodes a protein similar to human NF2/Merlin (43% identity), and contains <u>F</u>our-Point-One <u>E</u>zrin <u>R</u>adixin and <u>M</u>oesin (FERM) N, B, and C domains at the N-terminus (Figure 2B). The *lq132* splice donor mutation occurred after the conserved FERM domain regions, and the *ok754* in-frame deletion removes the entire FERM C domain, including the putative actin-binding site (Figure 2B). RNAi of *nfm-1* caused embryonic lethality (SKOP *et al.* 2004). Thus, both *lq132* and *ok754* are likely not null alleles and retain some function. AQR migration defects in *ok754* were significantly strong than *lq132*, suggesting that *ok754* is a stronger allele than *lq132*.

### Mosaic analysis indicates a non-autonomous requirement for *nfm-1* in anterior AQR migration

Genetic mosaic analysis using a rescuing *nfm-1(+)* extrachromosomal array was used to test if *nfm-1* was required in the Q cells themselves (see Methods). In *C. elegans*, extrachromosomal arrays are not stably inherited meiotically, and can be lost during mitotic cell divisions, creating genetically mosaic animals. We used an established strategy to score mosaic animals that had lost an *nfm-1(+)* rescuing transgene in AQR or PQR lineage (see Methods and (CHAPMAN *et al.* 2008; SUNDARARAJAN *et al.* 2014)). This strategy uses a stable *Pgcy-32::cfp* integrated transgene to visualize AQR and PQR in all animals, and an unstable array *nfm-1(+)*, carrying the rescuing *nfm-1::gfp* fosmid and *Pgcy-32::yfp* (Figure 4.3). Non-mosaic *nfm-1(ok754)* animals that harbored the *nfm-1(+)* array were viable, fertile, and were rescued for AQR and PQR migration (Figure 2E and Figure 2F). We analyzed 89 mosaic animals in which the *nfm-1(+)* array was lost from the AQR lineage, but retained in PQR and URX lineages as shown in Figure 3. These animals were rescued for AQR migration defects despite loss of *nfm-1(+)* in AQR, suggesting that *nfm-1* is required non-autonomously for anterior AQR migration (Figures 3B and Figures 3C and Figures 4A). Similarly, PQR migration defects were still rescued in 75 mosaic animals in which PQR that had lost the *nfm-1(+)* array (Figure 4B). Animals mosaic for *nfm-1(+)* in AQR or PQR rescued *nfm-1(ok754)* defects to a similar level as non-mosaics *(nfm-1(+)* in AQR and PQR) (Figure 4A and Figure 4B). In sum, this mosaic analysis indicates that *nfm-1* function was not required in AQR and PQR for their proper migration, and that *nfm-1* might act non-autonomously in this process.

**Figure 3.**
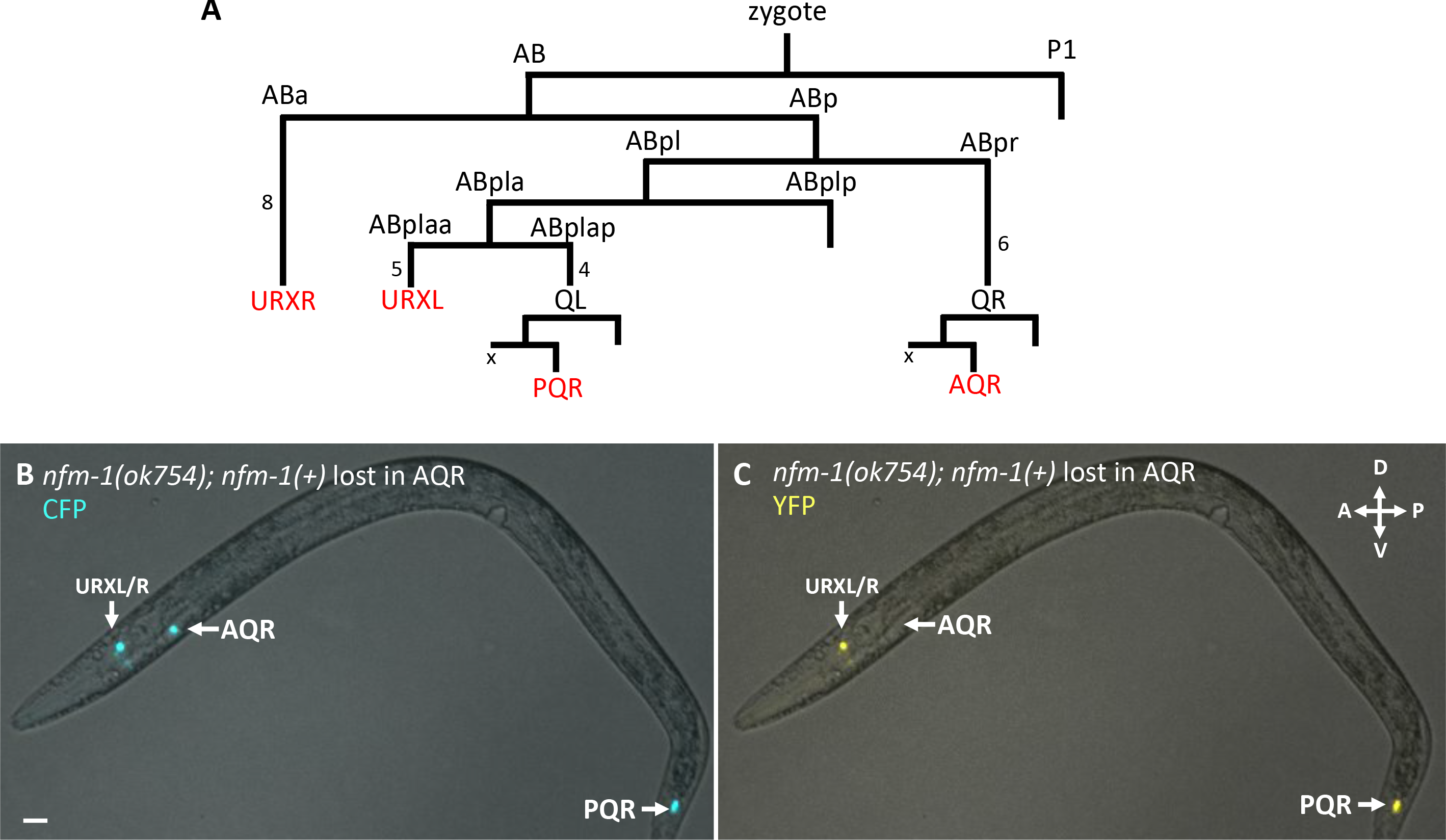
*nfm-1* mosaic analysis. A) The abbreviated lineage of cells that express *Pgcy-32* (red). Numbers next to lines indicate the number of cell divisions not shown. The X next to AQR and PQR indicates the sister of A/PQR (QL/R.aa) that undergoes programmed cell death. B) Fluorescent micrograph taken with CFP filter of *nfm-1(ok754); nfm-1(+), Pgcy-32::cfp* mosaic animal with correct placement of AQR and PQR. C) Fluorescent micrograph of the same animal from B using a YFP filter. AQR is not visible in this animal indicating that somewhere in AQR lineage, the *nfm-1(+)* transgene was lost. YFP is detected in URXL/R, and PQR indicating many tissues retained *nfm-1(+)*. Scale bar represent 10μM.

**Figure 4.**
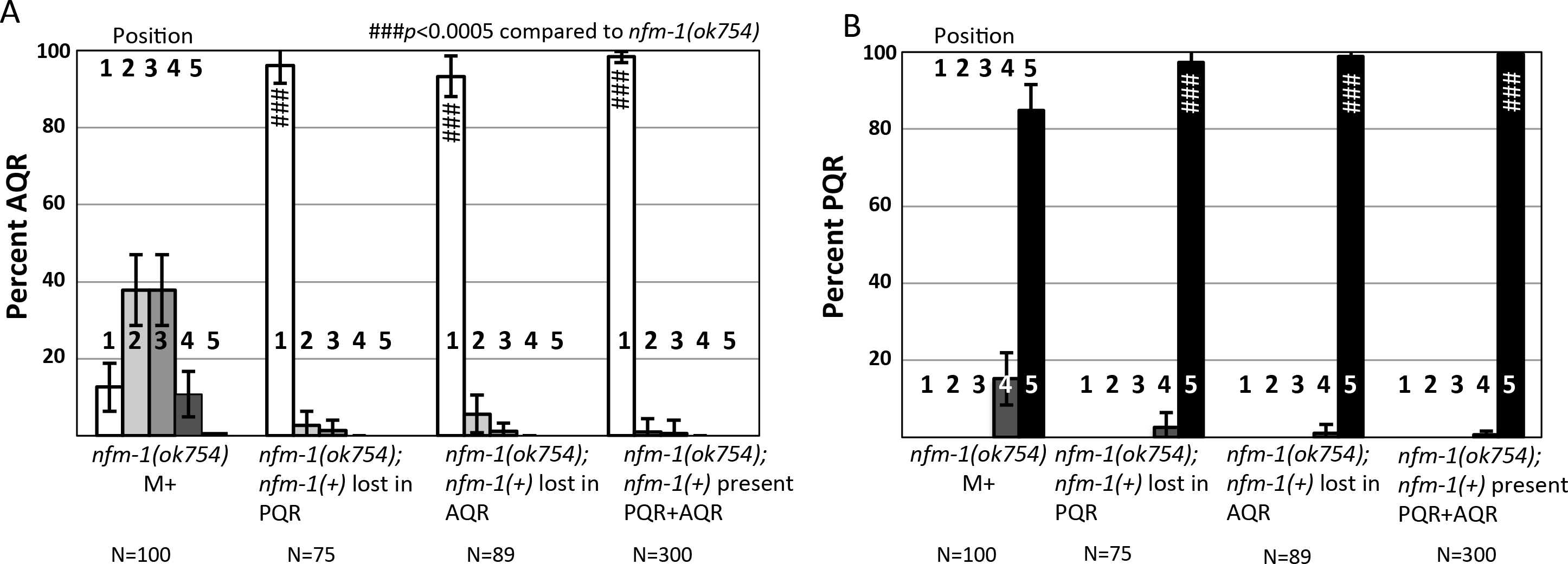
Analysis of *nfm-1(+)* mosaic animals. Quantification of AQR (A), and PQR (B) migration as in Figure 2, with *nfm-1(+)* mosaic animals. *nfm-1(+)* represents presence of *nfm-1* rescuing fosmid. *nfm-1(+)* rescued *ok754* lethality, and animals were maintained as rescued homozygous *ok754* mutants. Mosaic animals have *nfm-1(+)* in URX but have lost *nfm-1(+)* in either AQR or PQR. Pound signs indicate, for that position, a significant rescue of corresponding *nfm-1* mutant (N>100 #*p*<0.05, ##*p*<0.005, ###*p*<0.0005, Fisher’s exact test). Error bars represent two times the standard error of the proportion.

Using this method, it was impossible to determine which tissues require *nfm-1.* Based on cell lineages (Figure 3A), presence of *nfm-1(+)* array in URX and either AQR or PQR suggests it is present broadly in AB-derived lineages. P lineages were not assayed in this study, but mosaic *nfm-1(ok754)* animals, which normally arrest as sterile larvae, grew to viable and fertile adults, again suggesting broad *nfm-1* distribution in the mosaics, possibly including P-derived germ line and gonad lineages. it is possible that perdurance of NFM-1 protein, or array loss in the Q lineages themselves led to *nfm-1* function in the Q lineages despite loss in AQR or PQR. To account for these rare but possible events, we scored at least 70 mosaic animals. Overall, mosaic analysis suggests that *nfm-1* acts non-autonomously in AQR and PQR migration, as loss of the rescuing array in AQR or PQR did not correlate with mutant phenotype.

### *nfm-1::gfp* transcriptional and translational reporter expression was not apparent in Q lineages

A *Pnfm-1::gfp* transcriptional reporter was created by using a 2.1-kb region upstream of *nfm-1* to drive expression of GFP. This 2.1-kb region was the entire upstream region between *nfm-1* and the next gene *anmt-2.* At the time of Q migration, this construct showed expression in posterior cells near the anus, including posterior intestinal cells, the three rectal gland cells, and other unidentified cells that might be the anal sphincter muscle and the stomatointestinal muscle (Figure 5A–Figure 5 C). Variable expression in the hypodermis was also observed (Figure 5 A–Figure 5 C). Overlap between *Pnjm-1::gjp* and *Pegl-17::mCherry,* a Q cell marker, was not observed, consistent with *nfm-1* not being expressed in the Q neuroblasts during their migrations (Figure 5A–Figure 5C). Likewise, full length NFM-1::GFP expression from the rescuing fosmid was not observed in migrating Q cells (Figure 6). NFM-1::GFP was detected in the posterior gut region. In sum, no *nfm-1* transgene expression was observed in the Q cells during their migration.

**Figure 5.**
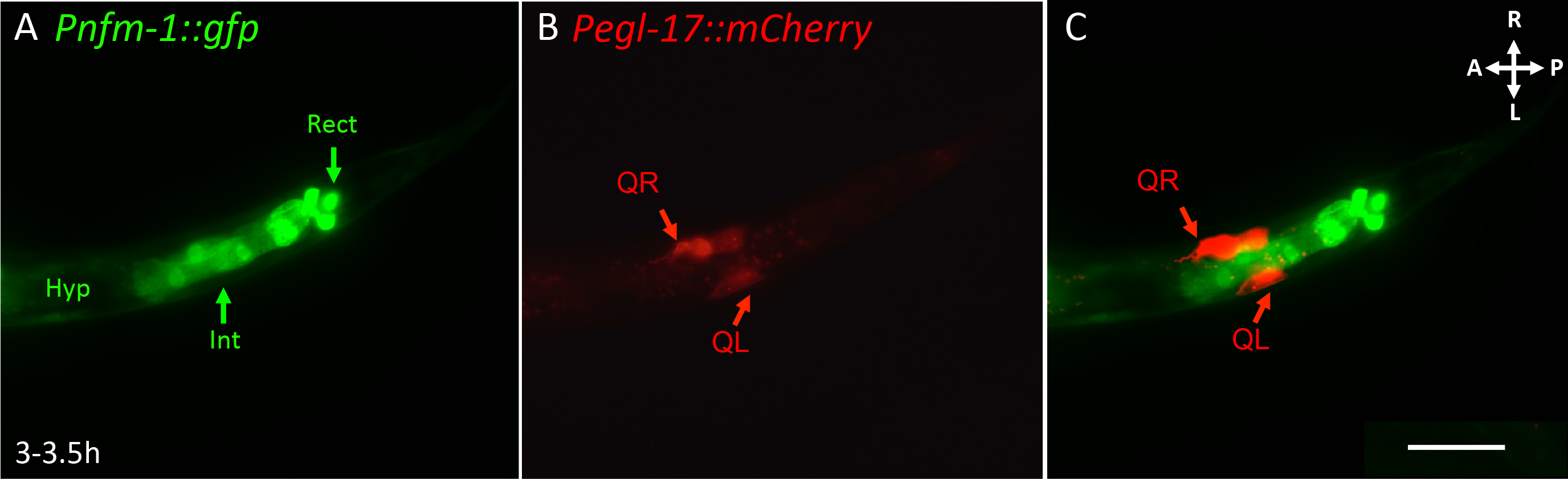
*nfm-1* transcriptional reporter was not expressed in Q cells during their early migrations. A-C) Ventral view of the posterior region of a *Pnfm-1::gfp; Pegl-17::mCherry* transgenic animal staged to 3-3.5h post hatching. A) A GFP Micrograph showing expression of *Pnfm-1::gfp.* Expression was seen in posterior cells near the anus, including posterior intestinal cells (Int) the three rectal gland cells (Rect). Other unidentified cells in the region were possibly the anal sphincter muscle and the stomatointestinal muscle. Variable hypodermal expression was observed along the length of the animal (Hyp). B) An mCherry micrograph shows Q cell specific expression during their migrations. C) Merged. GFP is not observed in Q cells, but is expressed in neighboring tissues and posterior cells. Scale bar is 10p,m, anterior is to the left, right is up.

**Figure 6.**
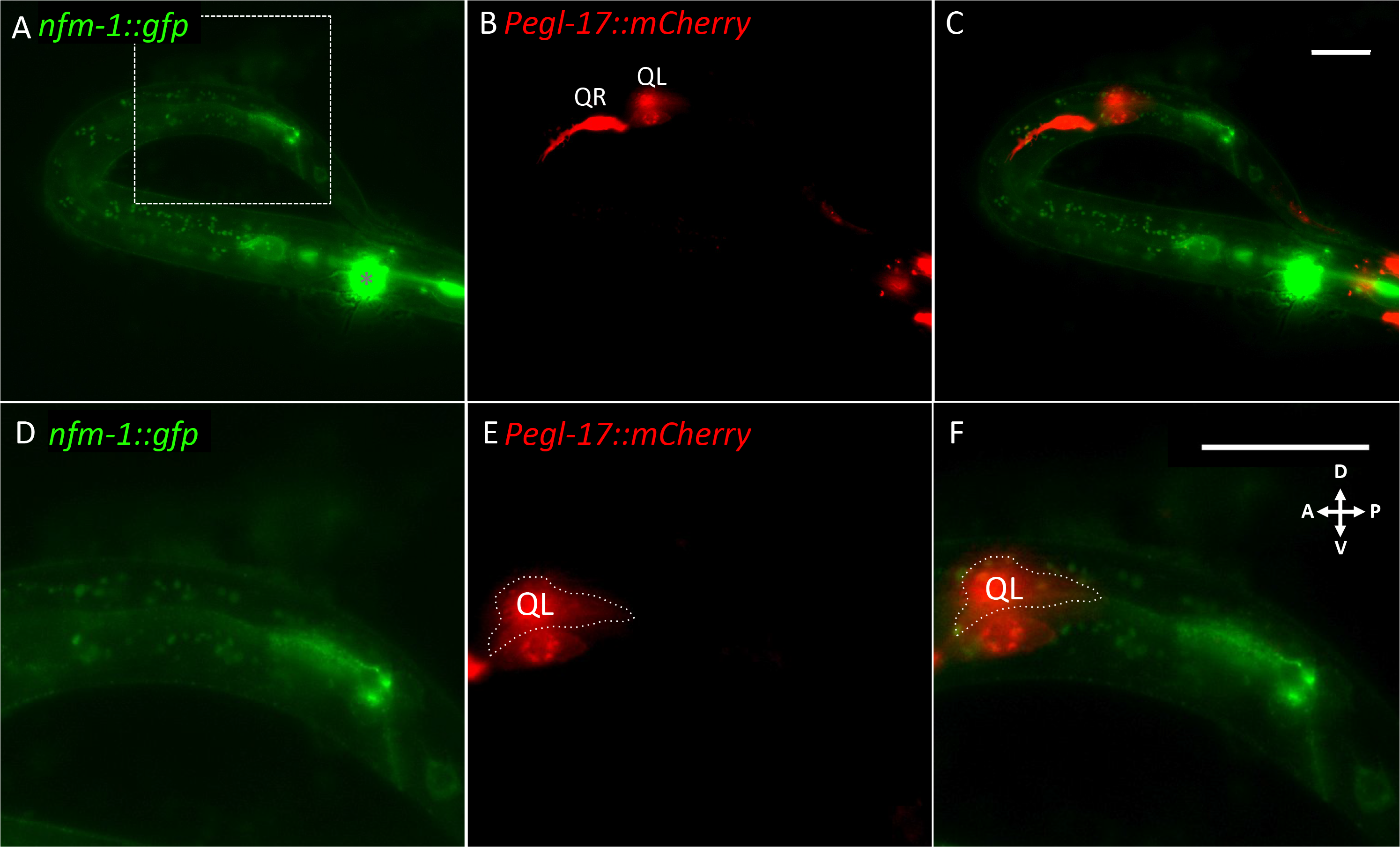
*nfm-1::gfp* translational reporter was excluded from Q cells. A-C) Lateral view of a staged 3-3-5h post hatching L1 *nfm-1::gfp; Pegl-17::mCherry* animal. A) Fluorescent micrograph of GFP expression from *nfm-1::gjp* rescuing fosmid. Asterisk marks URX expression of *Pgcy-32::yfp* in the head that was not excluded by GFP filter. The dashed rectangle indicates the enlarged posterior section in D-F. B) *Pegl-17::mCherry*, fluorescent micrograph showing location of early Q neuroblasts. QL is out of focus because QL and QR are on different planes, QR on the right side, and QL on left side of the animal. C) Merge of A and B. No overlap of mCherry and GFP was observed. D-F) Enlarged posterior section of A-C. Anterior is left, dorsal is up. D) Enlargement of A to show *nfm-1::gjp* present in posterior region near the anus. E) Enlargement of B. QL is outlined to distinguish it from the V5 seam cell that transiently expresses *Pegl-17*. F) Enlargement of C. Scale bars in C and F represent 10 μm.

### *slt-1* mutations enhance AQR defects of *nfm-1(lq132).*

Previous studies suggested that *NF2* can non-autonomously affect axon guidance in the developing mouse brain (LAVADO *et al.* 2014). This guidance mechanism occurs through regulation of *Slit2* mRNA levels, suggesting a transcriptional role of *NF2* (LAVADO *et al.* 2014). *Slit2* is a secreted guidance cue for developing neurons, and is detected by the Robo receptor. Because of interactions between *Slit2* and *NF2* we investigated the interaction of *nfm-1* and the *C. elegans Slit2* homolog *slt-1* in Q descendant migration. In this study we used one null allele *slt-1(eh15)*, and one strong loss-of-function in frame deletion allele *slt-1(ok255)* (HAO *et al.* 2001; STEIMEL *et al.* 2013). *slt-1* mutations had no effect on AQR and PQR migration on their own, but enhanced AQR migration defects of *nfm-1(lq132)* and *nfm-1(ok754)* (Figure 7). *slt-1* had no effect on PQR migration in double mutants. We tested the SLT-1 receptor SAX-3/Robo, and *sax-3(ky123)* mutants showed weak but significant defects in both AQR and PQR migration, consistent with SAX-3 promoting migration of the Q lineages (Figure 7).

**Figure 7.**
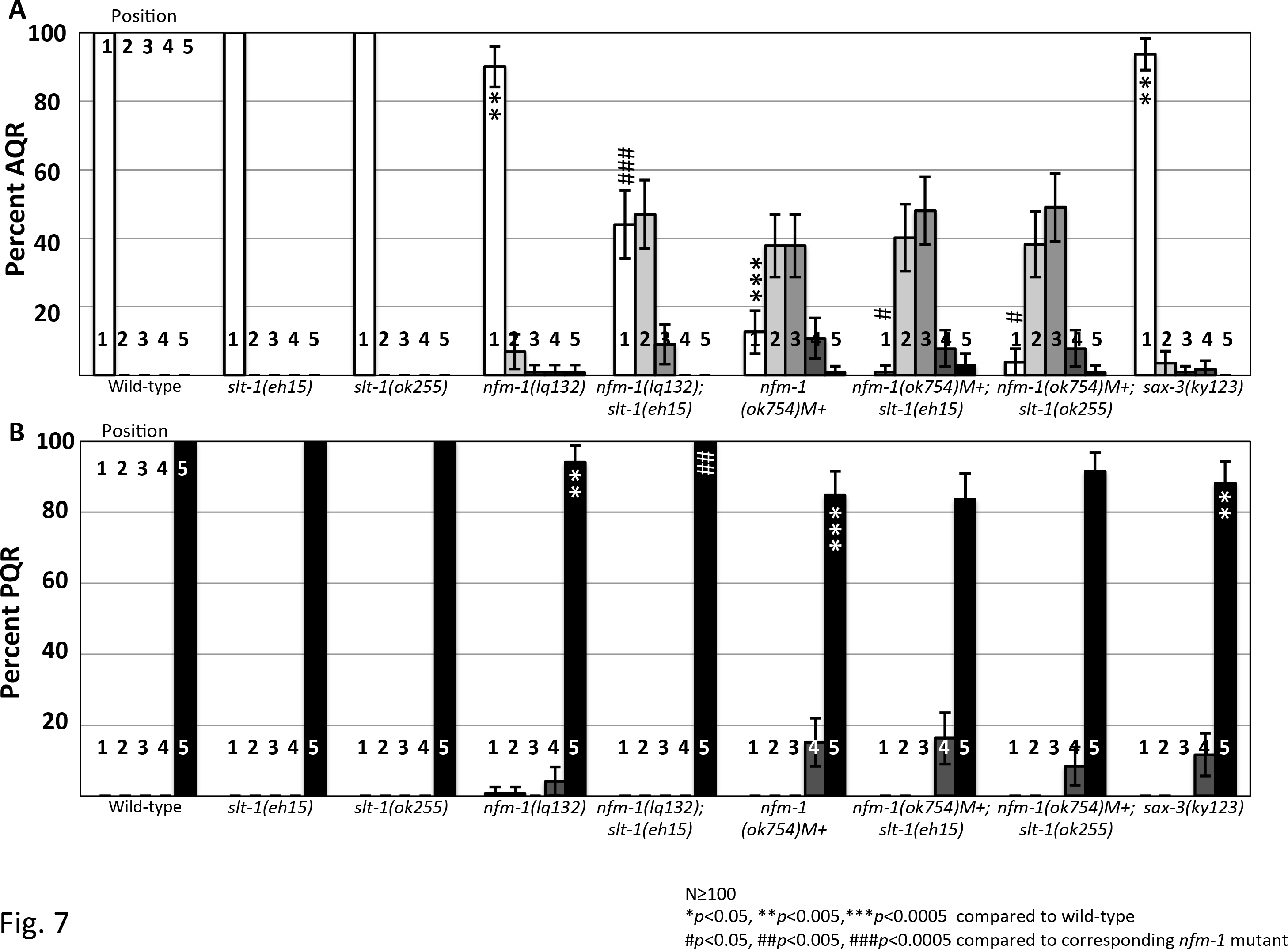
*slt-1* enhances *nfm-1* AQR migration defects. A) Percentage of AQR in each position, quantified as in Figure 2. B) PQR migration. Asterisks indicate significant difference from wild-type (N>100 \**p*<0.05, \*\**p*<0.005, \*\*\**p*<0.0005 Fisher’s exact test). Pound signs indicate, for that position, a significant enhancement of the corresponding *nfm-1* mutant (N>100 #*p*<0.05, ##*p*<0.005, ###*p*<0.0005, Fisher’s exact test). Error bars represent two times the standard error of the proportion.

## Discussion

### The NF2/Merlin molecule NFM-1 promotes migration of AQR and PQR

Complete migration of the QR and QL descendants AQR and PQR requires the coordination of many genes (MIDDELKOOP AND KORSWAGEN 2014). Although numerous molecules have been identified that act in the Q cells to promote migration, such as the transmembrane receptors UNC-40/DCC, PTP-3/LAR, and MIG-13 (SUNDARARAJAN AND LUNDQUIST 2012; WANG *et al.* 2013; SUNDARARAJAN *et al.* 2015), fewer have been identified that act outside the Q cells to control their migration. Of the non-autonomous genes that have been implicated in Q descendant migration, most are secreted molecules such as Wnts (HUNTER *et al.* 1999; WHANGBO AND KENYON 1999; KORSWAGEN 2002; PAN *et al.* 2006), although some transmembrane genes such as CDH-4 have been demonstrated to non-autonomously affect Q cell migration (SUNDARARAJAN *et al.* 2014).

Here we present data identifying a non-autonomous role for the FERM domain-containing molecule NFM-1 in promoting Q migration. NFM-1 is similar to human NF2/Merlin, the molecule affected in Neurofibromatosis type II. We found that mutations in *nfm-1* resulted in severe AQR migration defects, and to a lesser extent PQR migration defects. These defects typically manifested as incomplete migrations, suggesting that these *nfm-1* mutations did not affect direction of migration along the anterior/posterior axis, but rather the migratory capacity of these cells.

Loss of *NF2/Merlin* function in either mouse or *Drosophila* results in embryonic lethality (FEHON *et al.* 1997; MCCLATCHEY *et al.* 1997). In *C. elegans, nfm-1* appears to be required in embryonic development similar to other animals, as RNAi against *nfm-1* is reported as embryonic lethal (SKOP *et al.* 2004), and no null alleles of *nfm-1* have been described. The two *nfm-1* mutations studied here are not complete loss of function and likely retain some NFM-1 function. The 5’ splice site mutant *nfm-1(lq132)* was viable and fertile, and the in-frame deletion allele *nfm-1(ok754)* caused larval arrest. It is possible that complete loss of *nfm-1* function results in more severe AQR and PQR migration defects, possibly even directional defects, not observed in these alleles. The *nfm-1(ok754)* in-frame deletion removes part of the FERMB domain and the entire FERMC domain, suggesting that these domains are important in AQR and PQR migration.

### NFM-1 acts non-autonomously in AQR and PQR migration

As a cytoskeletal-membrane linker with a potential actin-binding domain, we hypothesized that NFM-1 might regulate actin-based membrane protrusion in migrating cells. However, a genetic mosaic analysis indicated that NFM-1 was not required in AQR or PQR for their migration. Furthermore, expression of transgenes of the *nfm-1* promoter and a rescuing full-length *nfm-1::gfp* were not observed in Q lineages. Rather, expression was observed in in posterior region near the anus, including posterior intestine, the rectal gland cells and potentially the anal sphincter muscle and stomatointestinal muscle. While the mosaic analysis was unable to discern the tissue in which NFM-1 acts, these expression studies suggest that NFM-1 function in the posterior region of the gut near the anus might regulate AQR and PQR migration. However, hypodermal expression is also a possibility. In sum, mosaic analysis and expression studies indicate that NFM-1 acts outside of the AQR and PQR, non-autonomously, to regulate their migration.

### *nfm-1* and *slt-1* interact genetically to promote anterior AQR migration

In *Drosophila* and mice, *NF2/Merlin* is known to regulate several signaling pathways, including stimulating the Hippo pathway to inhibit the Yorkie transcription cofactor (HAMARATOGLU *et al.* 2006; MOROISHI *et al.* 2015). In mice, loss of *NF2* in neural progenitor cells results in upregulation of Yap (LAVADO *et al.* 2014). High Yap activity leads to ectopic levels of the secreted guidance cue *Slit2* which causes defects in midline axon guidance (LAVADO *et al.* 2014). Interestingly, this is a non-autonomous role of *NF2* in midline axon guidance, similar to our observation of *nfm-1* in *C. elegans* neuronal migration. The Hippo pathway in *C. elegans* is poorly conserved, and the *C. elegans* genome does not encode a clear Yap homolog (HILMAN AND GAT 2011). We tested the role of the single *C. elegans Slit* gene *slt-1* in AQR/PQR migration and interaction with *nfm-1. slt-1* regulates the anterior-posterior migration of the CAN neurons in embryos (HAO *et al.* 2001). Although no migration defects were detected in *slt-1* mutants alone, they did enhance AQR migration defects of *nfm-1(lq132)* and *nfm-1(ok754)*. This enhancement is consistent with NFM-1 and SLT-1 acting in parallel pathways, but since we do not know the null phenotype of NFM-1 with regard to AQR and PQR, the possibility that they act in the same pathway cannot be excluded.

Interestingly no enhancement of *nfm-1* PQR migration defects was seen in *slt-1; nfm-1* double mutants. This suggests that *slt-1* and *nfm-1* interact in AQR migration but not PQR migration. This is of note because *nfm-1* is expressed in posterior tissues where PQR migrates, and away from which AQR migrates. *sax-3/Robo* mutants displayed both AQR and PQR migration defects. Possibly, SAX-3/Robo acts with SLT-1 in AQR migration, and with an unidentified ligand in PQR migration. In mice, midline axon defects are due to excess Slit2 expression in *NF2* mutants. The phenotypic enhancement that we observe between *slt-1* and *nfm-1* suggests that these molecules are both required for AQR migration. Further studies of the interaction between *nfm-1* and *slt-1* will be required to understand the role of these molecules in AQR migration.

Our results, combined with those in vertebrates, are consistent with the idea that NFM-1 promotes the production of a signal or signals that regulate AQR and PQR migration. This could be SLT-1 itself, such as in vertebrates, or a molecule that acts in parallel to SLT-1. The *slt-1* expression pattern is dynamic throughout development, but *slt-1* is expressed in posterior cells including body wall muscles and the anal sphincter muscle (HAO *et al.* 2001). Whether *nfm-1* can control pathways that regulate transcription as it does in vertebrates, or maintains epidermal integrity to control migration, is unclear, but our studies suggest that NFM-1 interacts with cues that guide AQR and PQR migrations.

## Acknowledgments

The authors thank members of the Lundquist and Ackley labs for discussion, and E. Struckhoff for technical assistance. Some *C. elegans* strains were provided by the CGC, which is funded by NIH Office of Research Infrastructure Programs (P40 0D010440). Some next-generation sequencing was provided by the University of Kansas Genome Sequencing Core Laboratory of the Center for Molecular Analysis of Disease Pathways (NIH P20 GM103638). This work was supported by NIH grants R01 NS040945 and R21 NS070417 to E.A.L., and the Kansas Infrastructure Network of Biomedical Research Excellence (NIH P20 GM103418). M.P.J was supported by the Madison and Lila Self Graduate Fellowship at the University of Kansas, and R.A. was a KINBRE Undergraduate Research Scholar (NIH P20 GM103418).

